# AP2 regulates Thickveins trafficking through Rab11 to attenuate NMJ growth signaling in *Drosophila*

**DOI:** 10.1101/2021.01.28.428584

**Authors:** Manish Kumar Dwivedi, Saumitra Dey Choudhury, Abhinandan Patnaik, Shirish Mishra, Raghu Padinjat, Vimlesh Kumar

**Author notes:** Present address: Section on Cellular Communication, National Institute of Child Health and Human Development, National Institutes of Health, 35 Convent Drive, Bethesda, MD 20892, USA. Equal contribution (Manish Kumar Dwivedi, Saumitra Dey Choudhury).

## Abstract

Compromised endocytosis in neurons leads to synapse overgrowth and altered organization of synaptic proteins. However, the molecular players and the signaling pathways which regulate the process remains poorly understood. Here we show that σ2-adaptin, one of the subunits of the AP2-complex, genetically interacts with BMP type I receptor, Thickveins (Tkv), and Daughter against decapentaplegic (Dad), two of the components of BMP signaling. We found that mutations in σ2-adaptin lead to an accumulation of Tkv receptors at the NMJ and results in a significant reduction in Tkv-positive early endosomes in the presynaptic terminals. Interestingly, the level of small GTPase Rab11 was significantly reduced in the σ2-adaptin mutant synapses. Consistent with the role of σ2-adaptin and Rab11 in the regulation of the same signaling pathway, a mutation in Rab11 or overexpression of a GDP-locked form of Rab11 (Rab11^S25N^) phenocopies the morphological and signaling defects of the σ2-adaptin mutants. Finally, we demonstrate that *σ2-adaptin* mutants show an accumulation of large vesicles and massive membranous structures, akin to endosomes at the synapse. Thus, we propose a model in which AP2 regulates Tkv internalization and recycling through a process that requires Rab11 activity to control the synaptic growth.

## INTRODUCTION

Synapse development and refinement is an interplay of signaling networks mediated by various endocytic, cytoskeletal, and actin regulatory proteins, Ubiquitin-Proteasome mediated protein degradation, Bone morphogenetic protein (BMP), and wingless (Wnt) pathways [1–8]. Understanding the crosstalk amongst them is crucial to our understanding of this process that regulate synapse development, refinement and plasticity [9]. BMP signaling pathway is a well-studied growth-promoting pathway at the *Drosophila* NMJ synapses [1, 7, 10, 11]. The canonical BMP signaling is dependent on phosphorylated Smad (pMAD in *Drosophila*) and its translocation in the ventral ganglion nuclei followed by the transcription of BMP target genes. At the *Drosophila* NMJ, the retrograde bone morphogenetic protein (BMP) signaling is initiated by secretion of glass bottom boat (Gbb) from the postsynaptic muscle. Gbb binds to wishful thinking (Wit, a type II receptor) and thickveins and saxophone (Tkv and Sax, type I receptors) at the presynaptic nerve terminals to control NMJ growth and function [7, 10, 12]. Gbb binding to Wit triggers the tetramerization of BMP receptors that, in turn, phosphorylates the Smad transcription factor, mothers against decapentaplegic (Mad). The phosphorylated form of Mad (pMAD) in complex with the co-Smad Medea is then retrogradely transported to the motor neuron nuclei, where it regulates gene transcription [13, 14].

Multiple studies have shown a tight correlation between defective endocytosis, altered synapse growth, and elevated synaptic phospho-MAD levels, which indicates increased BMP signaling [1, 8, 11, 15, 16]. One such study has shown that Nwk, an F-BAR and SH3 domain-containing protein that negatively regulates synaptic growth, interacts with Tkv along with Dap160 and Dynamin (both endocytic proteins) to attenuate retrograde BMP signaling during NMJ growth [11]. Endocytic and endosomal pathways are, therefore, critical to controlling both the activity and localization of signaling proteins that regulate synaptic growth [17, 18]. Clathrin-mediated endocytosis (CME) is required not only for basal synaptic transmission at nerve terminals but also for peripheral synapse development [3, 19, 20]. For instance, perturbations in CME resulting from mutations in Dynamin, AP2 subunits, Endo, or Synj all exhibit NMJ structural defects resulting in increased number but decreased size of synaptic boutons in *Drosophila* [3, 20]. Defects in intracellular trafficking can also lead to enhanced signaling from the cellular compartments (like endosomes) that has implications on synapse development [17, 21–23]. In the neuronal context, the efficacy of intercellular signaling is regulated by the trafficking of activated receptor/ligand complexes following endocytosis from the presynaptic membrane.

Tightly regulated endocytic transport of BMP receptors relies on the spatiotemporal regulation of Rab GTPase function [24]. The Rab-family of GTPases regulates the progression of receptor endocytosis and participates in the successive steps of membrane maturation, receptor transport, and turnover [25]. In particular, Rab5 regulates vesicle formation and is associated with early endosomes, while Rab7 and Rab11 associate with late and recycling endosomes, respectively [26, 27]. Endosomal trafficking of BMP signaling complexes at the nerve terminals is known to fine-tune the intensity and persistence of BMP signaling [9, 18]. Altered distribution or misregulation of Rab11 has been shown to suppress Tkv trafficking from early endosome to pre-synaptic membrane resulting in elevated BMP signaling [9, 11, 28, 29]. An important, yet enigmatic question, is the correlation between defective CME and aberrant synaptic growth. For instance, it is not known whether specific modes of endocytosis internalize specific cargos whose trafficking defect perturbs synaptic signaling. Similarly, it is unclear whether the NMJ structural defects associated with the endocytic mutants is a consequence of deficient endosomal trafficking leading to aberrant synaptic signaling. It is likely that perturbing CME deregulates signaling modules of BMP pathway that leads to elevated pMAD in the endocytic mutants [3, 11].

In central synapses, AP2-dependent CME is dispensable for membrane regeneration from the presynaptic plasma membrane following high-frequency nerve stimulation [30]; the critical role of CME in generating vesicles from endosome-like structures following bulk membrane endocytosis cannot be ruled out [31]. Previous studies support a model in which compromised CME can lead to defective signalosome trafficking by trapping signaling molecules in endosomes or intermediate structures of the endosomal pathway [22, 23, 32–35]. A recent study has highlighted the role of BMP receptor macropinocytosis to restrain BMP-mediated synaptic development by linking Abl and Rac1 GTPase signaling, indicating fine-tuning of endosomal trafficking of BMP receptors by small GTPases [36].

Our previous study has shown elevated levels of synaptic as well as motor-nuclei pMAD in *σ2-adaptin* mutants [3]. In order to investigate the underlying signaling mechanisms leading to elevated pMAD levels, we performed epistatic interactions between *σ2-adaptin* mutants with the components of BMP signaling. Our studies show that σ2-adaptin is required for internalization and endosomal trafficking of the BMP receptor Tkv at the NMJ synapses. Analysis of endocytic trafficking using endosomal markers suggests that defective Tkv receptor trafficking in *σ2-adaptin* mutants is a consequence of reduced synaptic Rab11 levels. Finally, our ultrastructural analysis of NMJ reveals the accumulation of large vesicles and supports a role of σ2-adaptin in the generation of signalosomes containing vesicles, possibly from endosomal structures. Thus, our studies reveal a novel function of σ2-adaptin in attenuating BMP-signaling by facilitating trafficking and recycling of the Tkv receptor through Rab11 containing recycling-endosomes.

## RESULTS

### *σ2-adaptin* genetically interacts with regulators of BMP signaling

In a previous study, we have shown that mutations in *σ2-adaptin* cause an increase in bouton numbers at NMJ as well as upregulation of pMAD, an effector of the BMP pathway [3]. To explore the role σ2-adaptin in regulating BMP signaling at the NMJ, we first assessed the epistatic interaction between *σ2-adaptin* and components of the BMP-signaling pathway. We found that introducing one mutant copy of the BMP type I receptor, Thickveins (Tkv) or co-Smad, Medea in *σ2-adaptin* mutant background could significantly suppress the synaptic overgrowth phenotype in these animals (Figure 1A-E). The number of boutons was significantly rescued in *tkv*^*7*^/+; *AP2σ*^*KG02457*^/*AP2σ*^*ang7*^: (1.91 ± 0.12, p ≤ 0.001) and *medea, AP2σ*^*KG02457*^/*AP2σ*^*ang7*^: (1.81 ± 0.07, p ≤ 0.001) when compared to *AP2σ*^*KG02457*^/ *AP2σ*^*ang7*^ (2.85 ± 0.08, p ≤ 0.001). However, there was no significant difference between wild-type control, heterozygous *AP2σ*^*KG02457*^/+, and *tkv*^*7*^/+ (WT: 1.26 ± 0.06, *AP2σ*^*KG02457*^/+: 1.33 ± 0.03 and *tkv*^*7*^/+: 1.34 ± 0.05) (Figure 1F).

**Figure 1.**
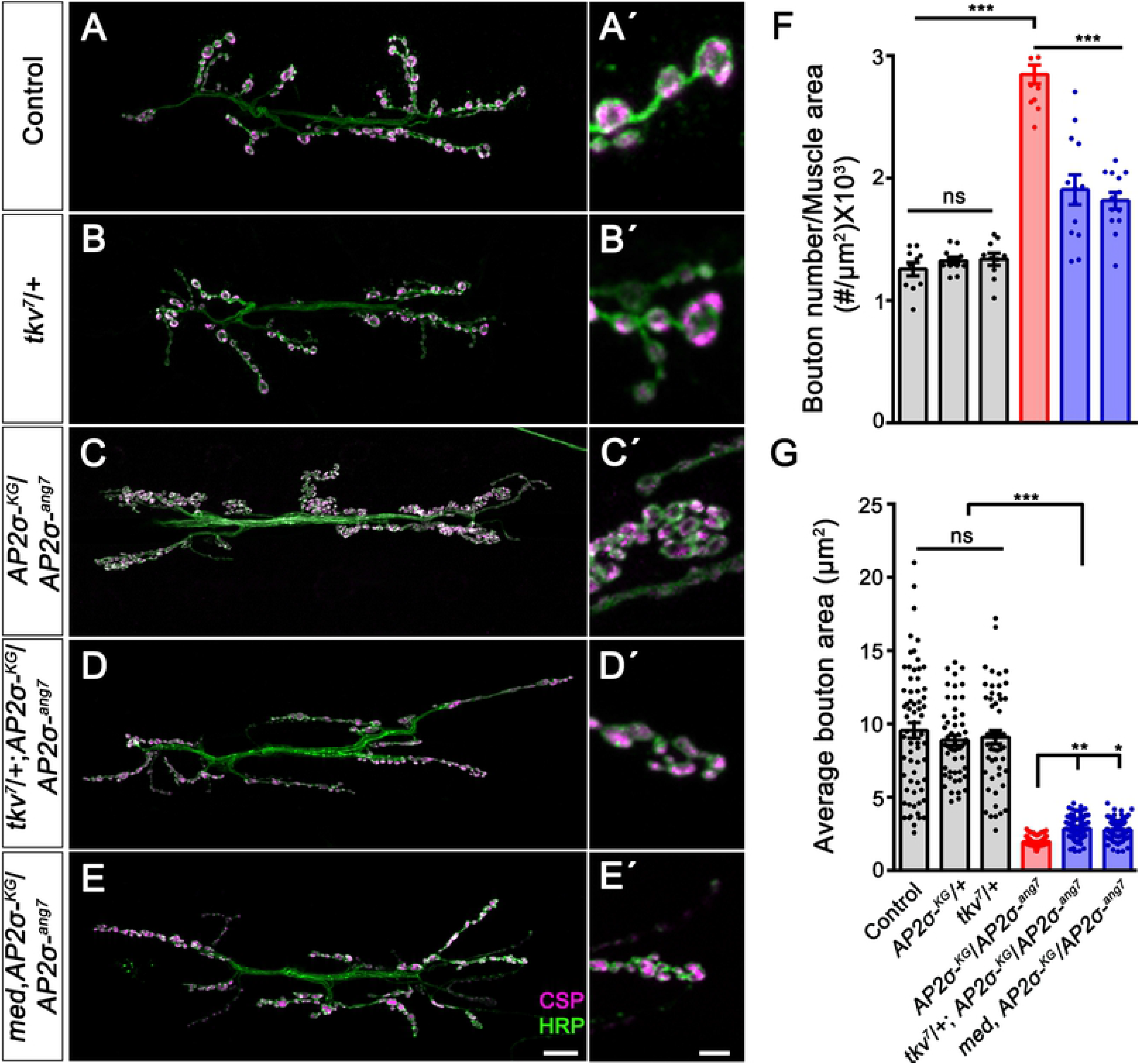
Downregulating BMP signaling components rescue the synaptic overgrowth and clustering in the *σ2-adaptin* mutant. (**A-E**’) Confocal images of NMJ synapses at muscles 6/7 of A2 hemisegment showing the synaptic growth in (A, A’) *w*^*1118*^ (Control), (B, B’) *tkv*^*7*^/+, (C, C’) *AP2σ*^*KG02457*^/*AP2σ*^*ang7*^, (D, D’) *tkv*^*7*^/+; *AP2σ*^*KG02457*^/*AP2σ*^*ang7*^, (E, E’) *medea*, *AP2σ*^*KG02457*^*/AP2σ*^*ang7*^ double immunolabeled with a pre-synaptic vesicle marker, CSP (magenta) and a neuronal membrane marker, HRP (green) to mark the bouton outline. Reducing the levels of BMP signaling components in the *AP2σ*^*KG02457*^*/AP2σ*^*ang7*^ background rescues the synaptic overgrowth. Scale bar in E (for A-E) and E’ (for A’-E’) represents 20 and 5 μm, respectively. (**F**) Histogram showing the average bouton number normalized to the muscle area from muscle 6/7 NMJ at A2 hemisegment in control animals (1.26 ± 0.06), *AP2σ*^*KG02457*^/+ (1.33 ± 0.03), *tkv*^*7*^/+ (1.34 ± 0.05), *AP2σ*^*KG02457*^*/AP2σ*^*ang7*^ (2.85 ± 0.08), *tkv*^*7*^/+; *AP2σ*^*KG02457*^*/AP2σ*^*ang7*^ (1.91 ± 0.12) and *medea*, *AP2σ*^*KG02457*^*/AP2σ*^*ang7*^ (1.82 ± 0.07). Error bar represents standard error of the mean (SEM); the statistical analysis was done using one-way ANOVA followed by post-hoc Tukey’s test. (**G**) Histogram showing the average bouton area from muscle 6/7 NMJ at A2 hemisegment in control animals (9.57 ± 0.53), *AP2σ*^*KG02457*^/+ (8.89 ± 0.35), *tkv*^*7*^/+ (9.11 ± 0.46), *AP2σ*^*KG02457*^/*AP2σ*^*ang7*^ (1.98 ± 0.03), *tkv*^*7*^/+; *AP2σ*^*KG02457*^/*AP2σ*^*ang7*^ (2.85 ± 0.05) and *medea*, *AP2σ*^*KG02457*^*/AP2σ*^*ang7*^ (2.79 ± 0.05). Error bar represents standard error of the mean (SEM); the statistical analysis was done using one-way ANOVA followed by post-hoc Tukey’s test. *p<0.05, **p<0.01, ***p<0.001; ns, not significant.

Consistent with the above observations, we found that mutating one copy of the type II BMP receptor, *wit* could also significantly rescue the synaptic overgrowth phenotype in *σ2-adaptin* mutants, (*wit*^*A12*^, *AP2σ*^*KG02457*^/*AP2σ*^*ang7*^: 2.00 ± 0.08 vs. *AP2σ*^*KG02457*^/*AP2σ*^*ang7*^: 2.50 ± 0.11; p ≤ 0.01) (Figure S1). Since elevated BMP signaling results in the formation of smaller boutons, we quantified the bouton area in these genotypes. We found that introducing one copy of *tkv*^*7*^ (*tkv*^*7*^/+; *AP2σ*^*KG02457*^/*AP2σ*^*ang7*^: 2.9 ± 0.05, p≤ 0.01) or *medea* (*medea, AP2σ*^*KG02457*^/*AP2σ*^*ang7*^: 2.79 ± 0.05, p ≤ 0.05) in *σ2-adaptin* mutant background slightly but significantly rescued the bouton area when compared to *σ2-adaptin* mutant alone (*AP2σ*^*KG02457*^/*AP2σ*^*ang7*^:1.97 ± 0.02) (Figure 1G). We found that partially downregulating these BMP pathway molecules reduces the clustering of boutons at the mutant NMJ (Figure 1A’-1E’). Thus, our data suggest that σ2-adaptin genetically interacts with the BMP receptors to regulate NMJ morphology.

In order to assess whether elevated BMP signaling indeed was responsible for the neuronal overgrowth in *σ2-adaptin* mutant, we tested the interaction between σ2-adaptin and the inhibitory Smad, Daughters against decapentaplegic (Dad), a negative regulator of BMP signaling [11, 37]. Phosphorylated MAD (pMAD) interacts with the co-Smad, Medea, and gets translocated to the nucleus to activate BMP target genes. Dad competes with Medea for binding with pMAD, which prevents its translocation into the nucleus and inhibits the relay of BMP signal [38]. *dad* loss-of-function mutant shows neuronal overgrowth with an increased number of satellite boutons, a phenotype that is strikingly similar to the endocytic mutants [11, 15]. We examined the total number of synaptic boutons in transheterozygotes of *σ2-adaptin* and *dad* mutants (Figure 2A-D). While the number of boutons in larvae heterozygous for *σ2-adaptin AP2σ*^*KG02457*^/+ (1.23 ± 0.05) and *dad*, *dad*^*j1E4*^/*+* (1.39 ± 0.09) was comparable with wild-type control (1.26 ± 0.06), transheterozygous *dad*^*j1E4*^/*AP2σ*^*KG02457*^ (1.89 ± 0.09, p≤0.001) showed significantly higher bouton number when compared to controls (Figure 2E). Taken together, our data indicate that σ2-adaptin may regulate BMP signaling at the *Drosophila* NMJ.

**Figure 2.**
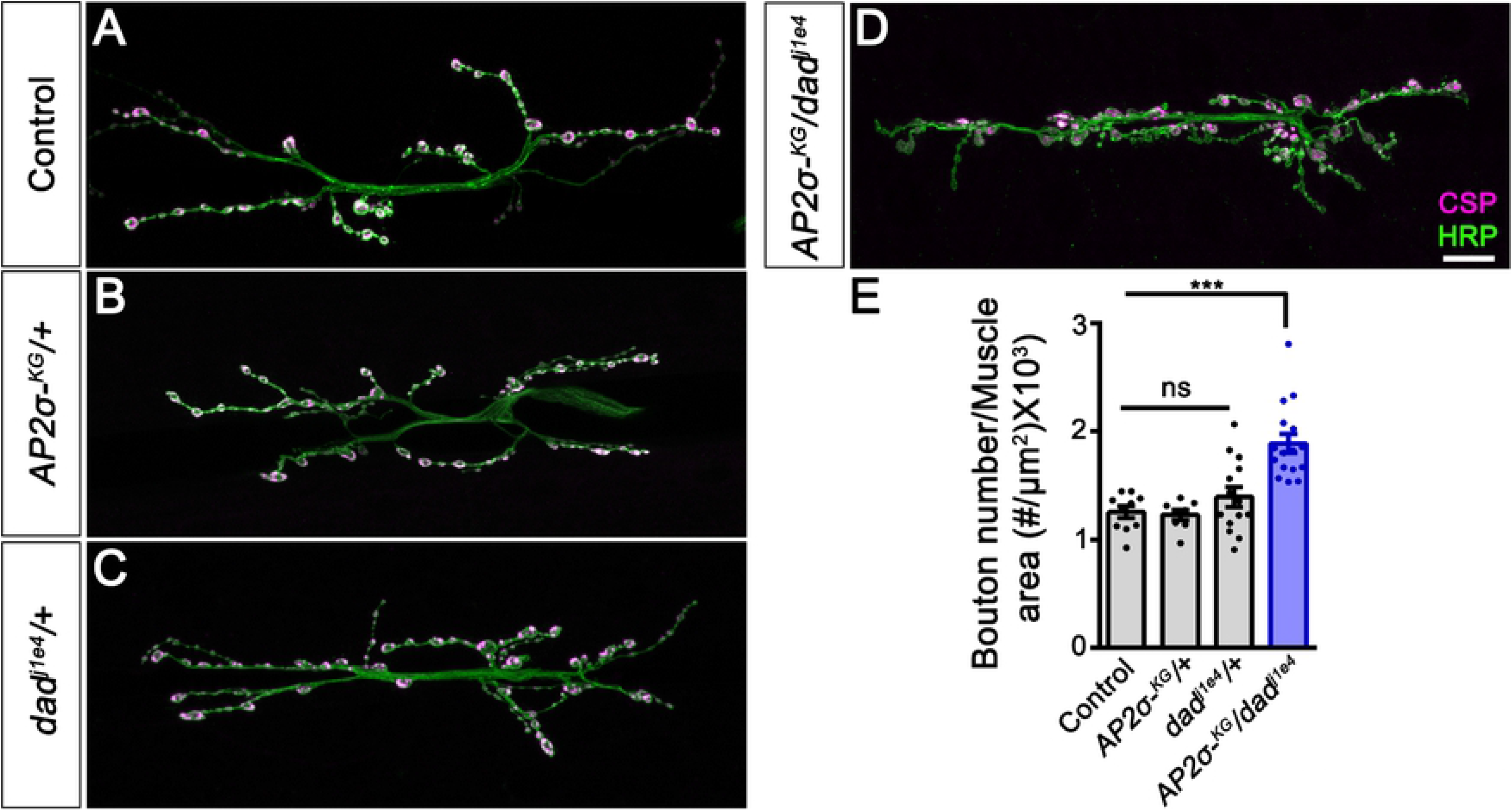
*σ2-adaptin* genetically interacts with *dad*, the inhibitory Smad of BMP signaling. (**A-D**) Confocal images of NMJ synapses at muscle 6/7 NMJ at A2 hemisegment showing the synaptic growth in (A) Control (*w*^*1118*^), (B) *AP2σ*^*KG02457*^/+, (C) *dad*^*j1E4*^/+, and (D) *AP2σ*^*KG02457*^/ *dad*^*j1E4*^ double immunolabeled with a pre-synaptic synaptic vesicle marker, CSP (magenta) and a neuronal membrane marker, HRP (green) to mark the bouton outline. *σ2-adaptin* and *dad* interact genetically, and trans-heterozygotes of *σ2-adaptin* and *dad* mutants show significantly increased synaptic growth. Scale bar in (D) represents 20 μm. (**E**) Histogram showing the average bouton number normalized to the muscle area from muscle 6/7 NMJ at A2 hemisegment in control animals (1.26 ± 0.06), *AP2σ*^*KG02457*^/+ (1.23 ± 0.05), *dad*^*j1E4*^/+ (1.39 ± 0.09) and *AP2σ*^*KG02457*^/ *dad*^*j1E4*^ (1.89 ± 0.09). Error bar represents standard error of the mean (SEM); the statistical analysis was done using one-way ANOVA followed by post-hoc Tukey’s test. \*\*\**p*<0.001; ns, not significant.

### The functional and morphological aspect of σ2-adaptin can be genetically delineated

Since one mutant copy of the Tkv receptor in the *σ2-adaptin* mutant background significantly restores the morphological defects, we asked whether the electrophysiological defects associated with the *σ2-adaptin* mutant are also rescued. We measured evoked excitatory junction potential (EJP), quantal content (QC), and high-frequency intracellular recording on wild-type, *AP2σ*^*KG02457*^*/AP2σ*^*ang7*^ and *tkv*^*7*^/+; *AP2σ*^*KG02457*^*/AP2σ*^*ang7*^ larvae. We found that miniature excitatory junction potential (mEJP) amplitude and frequency, EJP amplitude, or the activity-dependent decline in the EJP amplitude in *tkv*^*7*^/+; *AP2σ*^*KG02457*^*/AP2σ*^*ang7*^ larvae were not significantly different than the *σ2-adaptin* mutant (Figure 3A-C). Moreover, we found that *tkv*^*7*^/+; *AP2σ*^*KG02457*^*/AP2σ*^*ang7*^ animals do not show a significant change in the quantal content compared to the *σ2-adaptin* mutants (*tkv*^*7*^/+; *AP2σ*^*KG02457*^*/AP2σ*^*ang7*^, QC= 38.20 ± 3.09 vs. *AP2σ*^*KG02457*^*/AP2σ*^*ang7*^, QC= 39.29 ± 5.81) (Figure 3D). These data suggest that while reducing the level of BMP signaling by lowering Tkv receptors in *σ2-adaptin* mutant partially rescues the morphological defects, it does not restore the physiological deficiencies in *σ2-adaptin* mutants suggesting that functional and morphological defects in *σ2-adaptin* mutant are independent of one another.

**Figure 3.**
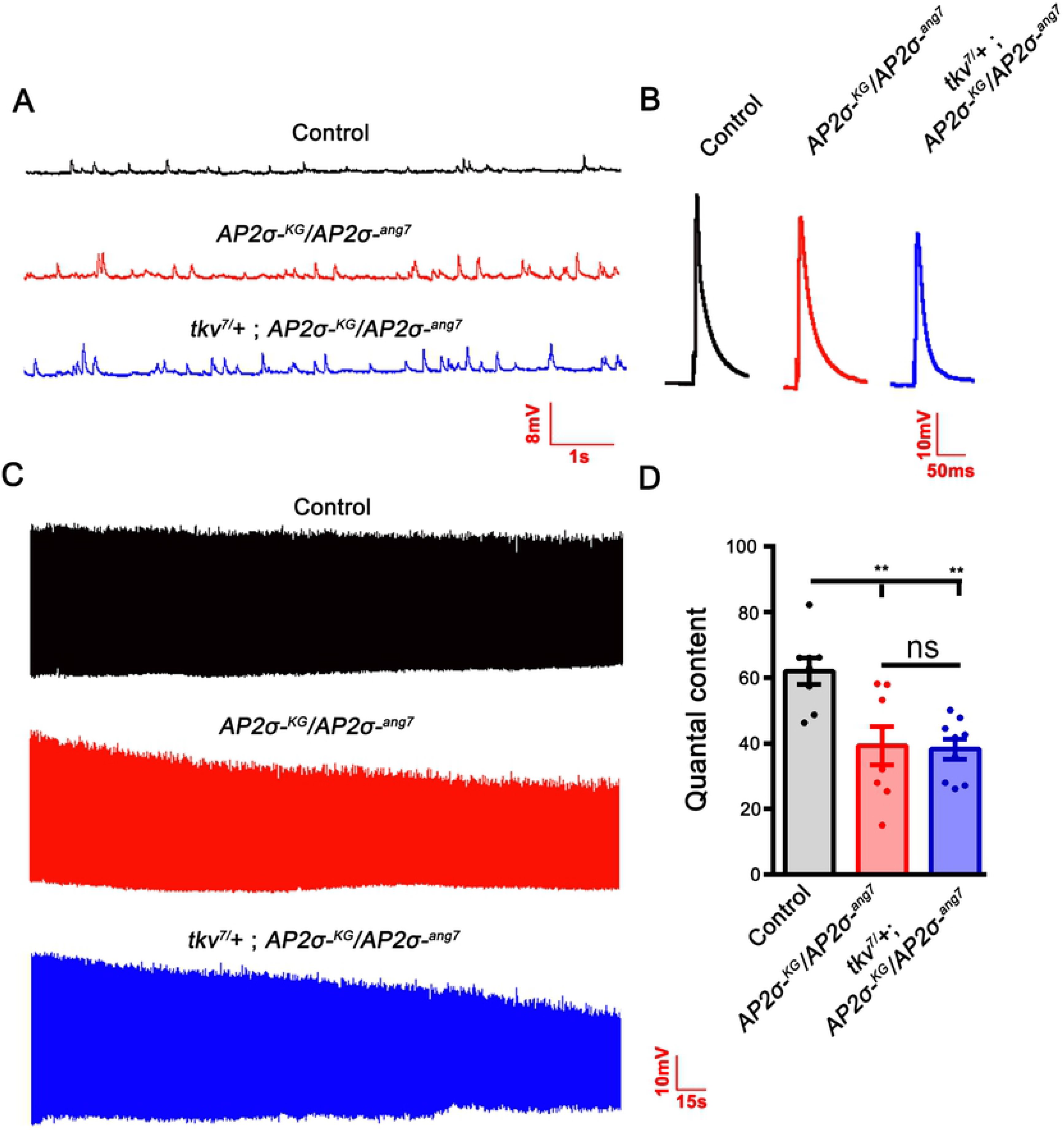
Structural and functional deficits in *σ2-adaptin* mutant can be genetically delineated. (**A**) Representative traces of mEJP in control, heteroallelic *AP2σ*^*KG02457*^*/AP2σ*^*ang7*^ and *tkv*^*7*^/+; *AP2σ*^*KG02457*^*/AP2σ*^*ang7*^ larvae. (**B**) Representative traces of EJP in control, heteroallelic *AP2σ*^*KG02457*^*/AP2σ*^*ang7*^ and *tkv*^*7*^/+; *AP2σ*^*KG02457*^*/AP2σ*^*ang7*^ larvae. (**C**) Representative traces of EJPs under high-frequency stimulation of control, heteroallelic *AP2σ*^*KG02457*^*/AP2σ*^*ang7*^ and *tkv*^*7*^/+; *AP2σ*^*KG02457*^*/AP2σ*^*ang7*^ larvae stimulated at 10 Hz for 5 min in 1.5 mM Ca^2+^ containing HL3. (**D**) Quantification of quantal content in control (61.92 ± 4.02), heteroallelic *AP2σ*^*KG02457*^/*AP2σ*^*ang7*^ (39.29 ± 5.81) and *tkv*^*7*^/+; *AP2σ*^*KG02457*^*/AP2σ*^*ang7*^ (38.2 ± 3.09). At least 8 NMJ recordings of each genotype were used for quantification. Error bars represent standard error of the mean (SEM); statistical analysis is based on one-way ANOVA followed by *post-hoc* Tukey’s multiple-comparison test. **p<0.01; ns, not significant.

### Loss of *σ2-adaptin* leads to the accumulation of endosome-like structures at the NMJ

Studies have shown that CME and Rab11 mediate the internalization and recycling of the BMP receptors [28, 39]. Altered levels of endosomal proteins such as Rab5 and Rab11 that are known to be involved in the trafficking of BMP receptors result in elevated BMP signaling leading to an increase in the number of boutons. We performed the NMJ transmission electron microscopy to understand how the loss of σ2-adaptin affects synapse ultrastructure. Interestingly, ultrastructural analysis of *σ2-adaptin* deficient synapses showed an accumulation of large endosome-like structures, similar to what has been shown in mutants that affect the endocytic and endosomal recycling machinery such as clathrin [40, 41], AP180 [42], Rab5 [43], Rab8 [44] and Rab11 [45]. We found a drastic decrease in the SV density (*w*^*1118*^: 85.18 ± 12.26 vs. *AP2σ*^*KG02457*^/ *AP2σ*^*ang7*^: 28.02 ± 14, p≤0.01) and an increase in the size of SVs (*w*^*1118*^: 43.16 ± 0.94 vs. *AP2σ*^*KG02457*^/ *AP2σ*^*ang7*^: 71.53 ± 3.7, p≤0.001) in the *σ2-adaptin* mutants. Moreover, we found large membrane invaginations in the mutant synapse, similar to what has been reported earlier for clathrin mutants [41]. These ultrastructural defects were rescued upon ubiquitous expression of a *σ2-adaptin* transgene (*actin5C*/+; *AP2σ*^*KG02457*^, *UAS-AP2σ/AP2σ*^*ang7*^: SV density (107.9 ± 11.32) and size (43.1 ± 1.46) (Figure 4A-E). Together, our data indicate that compromised regeneration of vesicles from the presynaptic membrane and defective membrane recycling in *σ2-adaptin* mutants results in the accumulation of large endosome-like structures.

**Figure 4.**
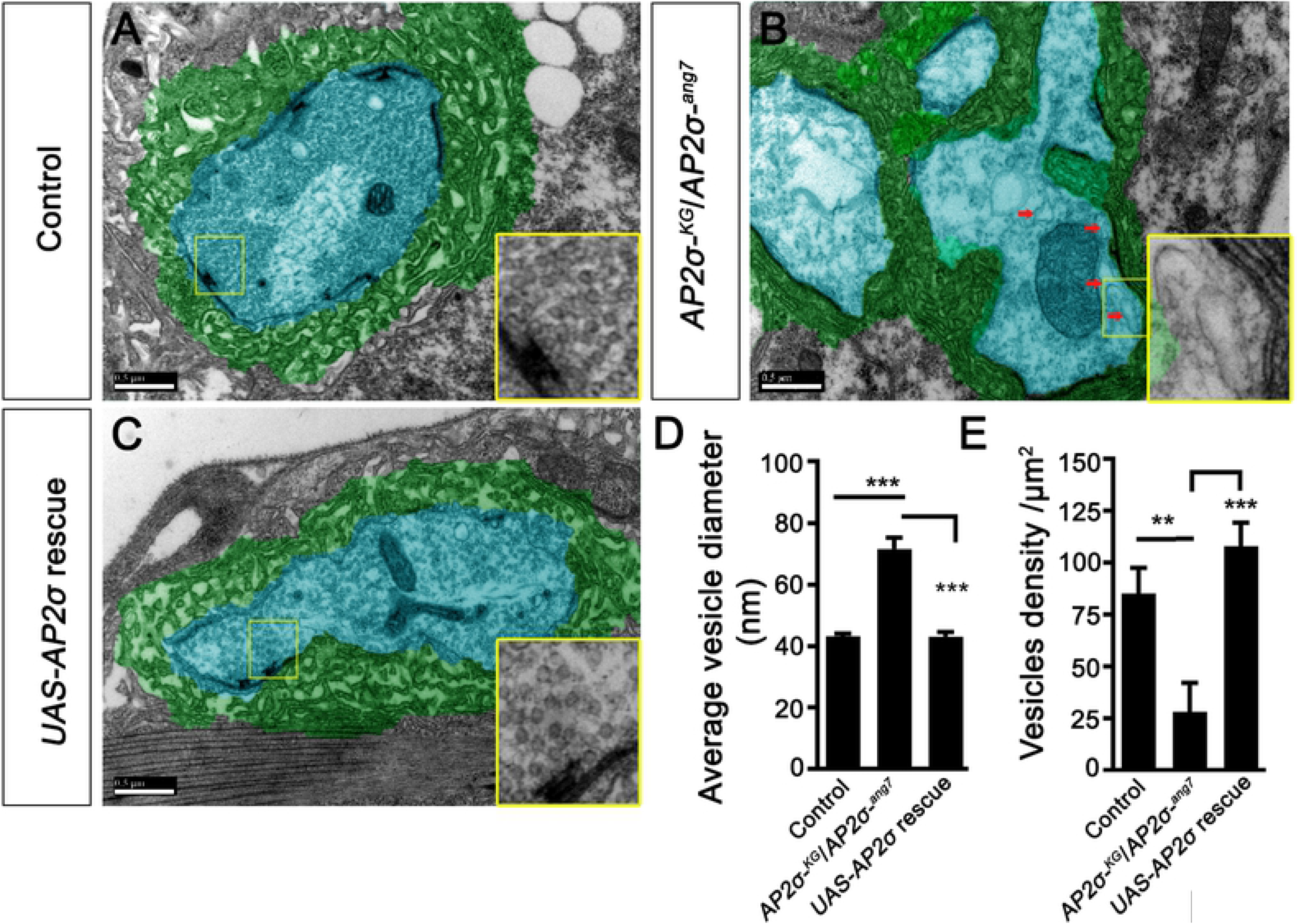
*σ2-adaptin* mutant synapses show accumulation of large endosomal structures. (**A-D**) Electron micrographs of third instar type Ib boutons of control (A), *AP2σ*^*KG02457*^*/AP2σ*^*ang7*^ (B), and *actin5C*/+; *AP2σ*^*KG02457*^, *UAS-AP2σ*/*AP2σ*^*ang7*^ (C). Arrows point to the large endosome-like structures observed in *AP2σ*^*KG02457*^*/AP2σ*^*ang7*^ boutons but are absent in control and rescued boutons. The insets show the magnified area around the active zones. The pre-synaptic compartment is pseudocolored in cyan, and the sub-synaptic reticulum is marked in green. Scale bar represents 500 nm. (**D**) Histogram showing average vesicle diameter in control (43.16 ± 0.94), *AP2σ*^*KG02457*^/*AP2σ*^*ang7*^ (71.53 ± 3.7), and *actin5C*/+; *AP2σ*^*KG02457*^, *UAS-AP2σ*/*AP2σ*^*ang7*^ (43.1 ± 1.46). (**E**) Histogram showing the SV density per unit area in control (85.18 ± 12.26), *AP2σ*^*KG02457*^*/AP2σ*^*ang7*^ (28.02 ± 14), and *actin5C*/+; *AP2σ*^*KG02457*^, *UAS-AP2σ*/*AP2σ*^*ang7*^ (107.9 ± 11.32). At least 10 images from three different larvae per genotype were used for quantification. Error bar represents standard error of the mean (SEM); the statistical analysis was done using one-way ANOVA followed by post-hoc Tukey’s test. \*\*\**p*<0.001, **p<0.01.

### *σ2-adaptin* mutants have increased synaptic Tkv receptors

AP2 complex has been shown to regulate CME and activity-dependent vesicle regeneration from endosome-like vacuoles [30, 46]. Because ultrastructural analysis of *σ2-adaptin* mutants revealed accumulation of membrane invaginations and large endosome-like structures similar to mutants with perturbed endosomal trafficking such as *Rab5*, *Rab11* and, *Rab8* mutants [43–45], we hypothesized that *σ2-adaptin* could be involved either in endocytosis of BMP receptors from the presynaptic membrane or in the endosome-dependent trafficking of the receptors. To check this possibility, we first assessed the level of Tkv receptors at the larval NMJ. Since a specific antibody against Tkv receptors is not available, we expressed an EGFP-tagged Tkv receptor transgene in the motor neurons of *σ2-adaptin* mutants (*D42-Gal4> UAS-Tkv-EGFP; AP2σ*^*ang7*^/*AP2σ-*^*KG02457*^). Interestingly, we found a significant accumulation of Tkv receptors at *σ2-adaptin* synapses (*D42-Gal4*, *AP2σ*^*KG02457*^/*AP2σ*^*ang7*^, *UAS-Tkv-EGFP*:135 ± 6.7, p≤0.01) when compared to control (*D42-Gal4*/*UAS-Tkv-EGFP*: 100 ± 9.36) (Figure 5A-E). This synaptic accumulation of Tkv-EGFP could be due to compromised endocytosis from the plasma membrane. To check this possibility, we analyzed the intensity profiles of Tkv receptor and the presynaptic membrane maker, HRP. When compared to the control synapses where Tkv localizes both at the presynaptic membrane as well as within the bouton, we found a higher intensity peak of Tkv-EGFP at the synaptic membrane of *σ2-adaptin* mutant (Figure 5F-K). Taken together, these data suggest endocytosis/trafficking of Tkv receptors in *σ2-adaptin* mutants are severely compromised, leading to its accumulation at the synaptic membranes.

**Figure 5.**
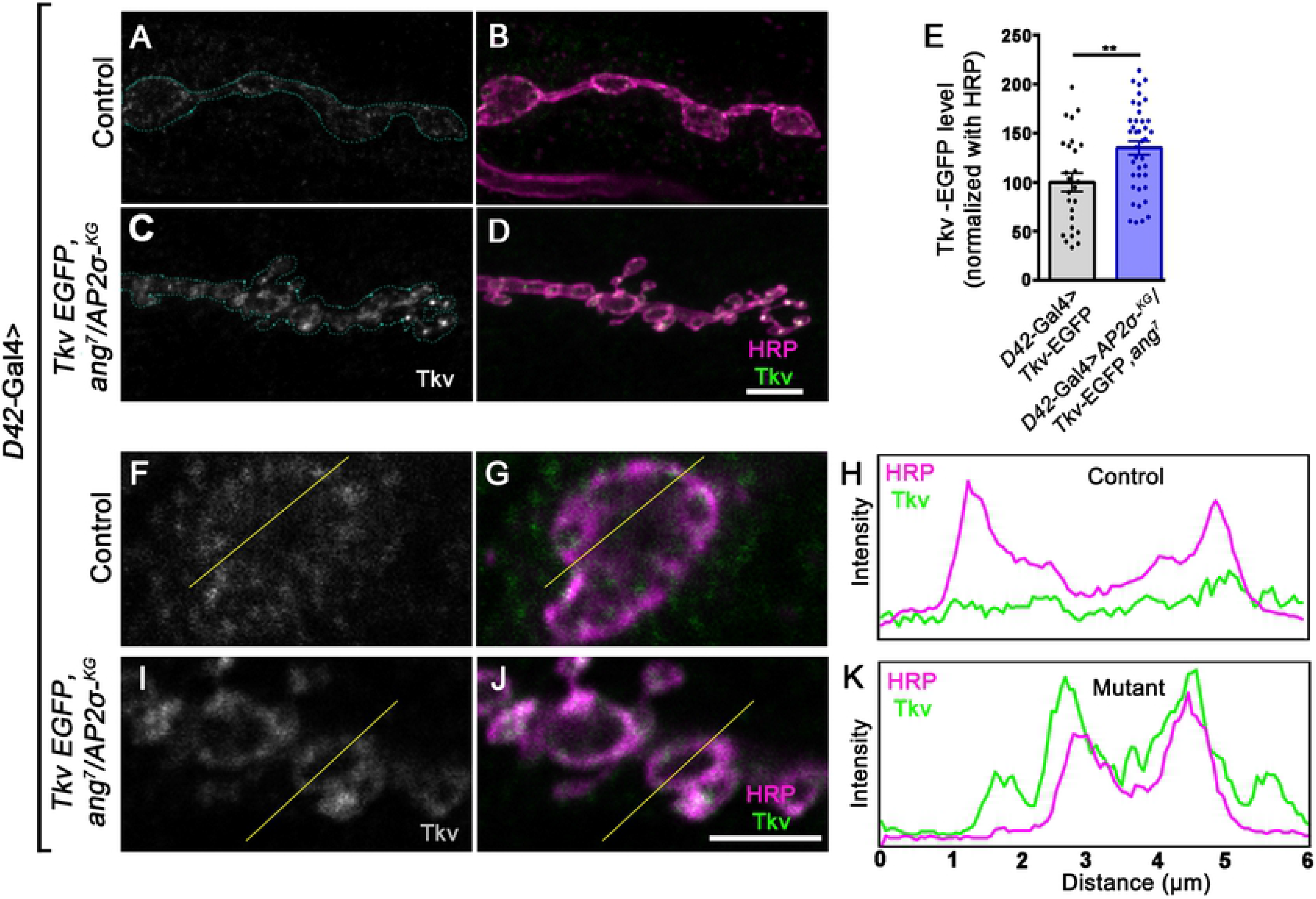
*σ2-adaptin* mutant synapses show accumulation of Tkv-EGFP. (**A-D**) Confocal images of NMJ synapses at muscle 4 NMJ at A2 hemisegment in *D42-Gal4*/*UAS-Tkv-EGFP* (A-B) and *D42-Gal4*, *AP2σ*^*KG02457*^/*AP2σ*^*ang7*^, *UAS-Tkv-EGFP* (C-D). The neuronal membrane is marked with HRP (magenta), and EGFP fluorescence is shown in greyscale. The bouton area is outlined in the grey channel. Scale bar in (D) represents 5 μm. (**E**) Histogram showing the relative Tkv level normalized to HRP in *D42-Gal4/UAS-Tkv-EGFP* (100 ± 9.36) and *D42-Gal4*, *AP2σ*^*KG02457*^/*AP2σ*^*ang7*^, *UAS-Tkv-EGFP* (135 ± 7) synapses. Error bar represents standard error of the mean (SEM); the statistical analysis was done using Student’s *t*-test. \*\**p*<0.01. (**F-K**) A single confocal section of a bouton labeled for Tkv (represented in grayscale) and presynaptic membrane marker HRP (magenta) in *D42-Gal4*/*UAS-Tkv-EGFP* (F-G) or *D42-Gal4*, *AP2σ*^*KG02457*^/*AP2σ*^*ang7*^, *UAS-Tkv-EGFP* (I-J). Note that the intensity profile plot across bouton (shown in G and J as a thin line) shows that compared to control (H), *σ2-adaptin* mutant (K) has more Tkv in at the membrane. Scale Bar in J represents 3 μm.

### *σ2-adaptin* mutation results in decreased levels of recycling endosomal marker Rab11

Receptors are known to be endocytosed, trafficked to the early endosomes, and sorted out for recycling or degradation [47–49]. Therefore, we hypothesized that the endocytosis of Tkv receptors from the presynaptic membrane should be compromised in the *σ2-adaptin* mutant. To test this prediction, we measured overall Tkv receptor levels at NMJ in *σ2-adaptin* mutants using Tkv-EGFP. We found a significant increase of Tkv receptors at NMJ in *σ2-adaptin* mutants. Further, we assessed the colocalization of Tkv-EGFP with an early endosomal marker, Rab5, at the NMJ. We found a drastic decrease in colocalization between Rab5 and Tkv-EGFP in *σ2-adaptin* mutant (*D42-Gal4*, *AP2σ*^*KG02457*^/*AP2σ*^*ang7*^, *UAS-Tkv-EGFP*: 27.40 ± 3.82, p≤0.001) when compared to control (*D42-Gal4/UAS-Tkv-EGFP*: 65.74 ± 3.75) (Figure 6A-G). Similar results were obtained when the same images were used to quantify the extent of colocalization using motion tracker software [50, 51]. Defective endosomal recycling results in the enrichment of activated receptors in early endosomes that cause an elevation in BMP signaling [28, 44, 52]. Enriched Tkv receptor levels at the synaptic membrane and its decreased colocalization with Rab5 in *σ2-adaptin* mutant suggests compromised internalization of Tkv receptor.

**Figure 6.**
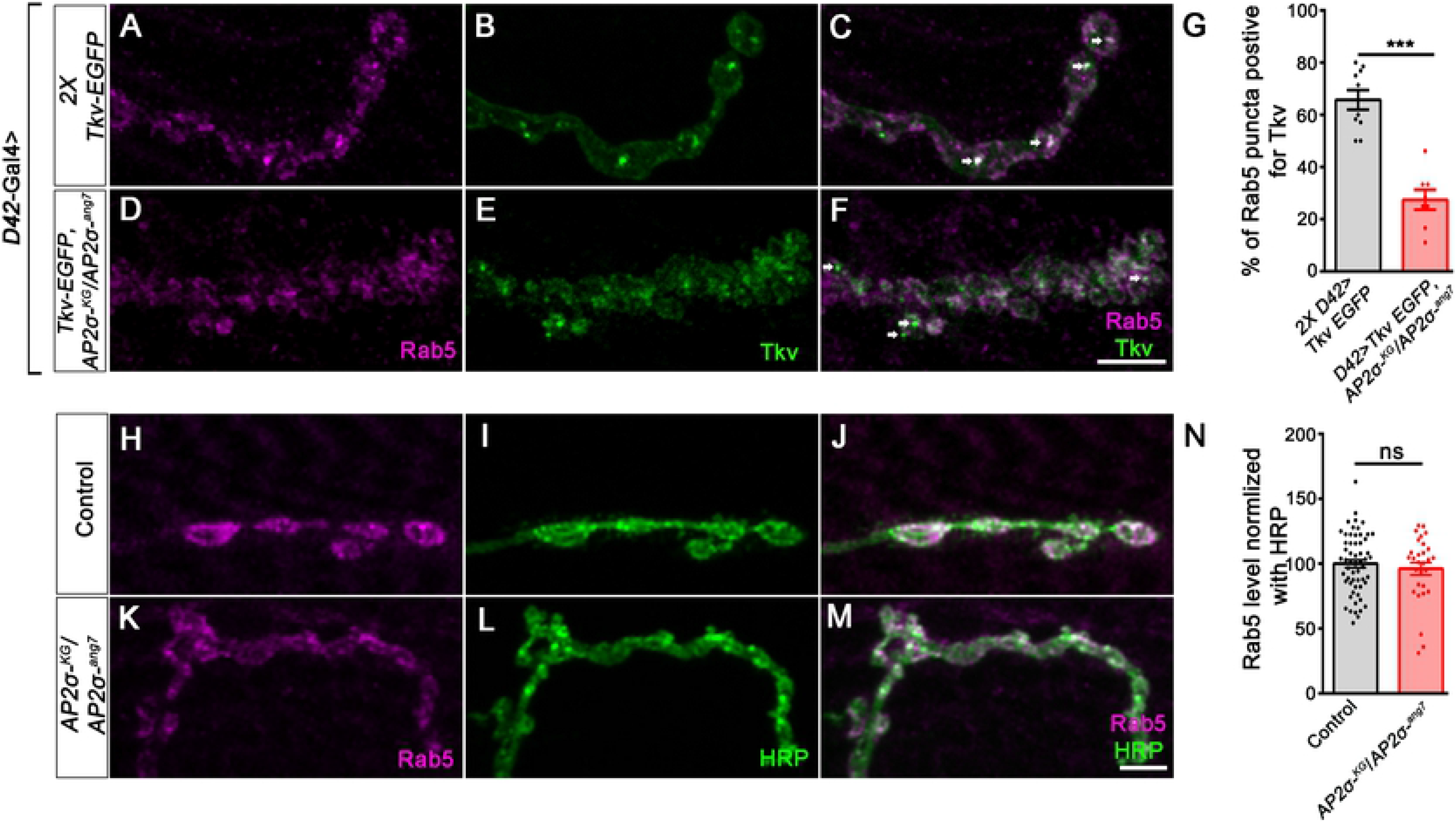
*σ2-adaptin* mutant synapses show reduced colocalization of Tkv-EGFP punctae with the early endosomal marker, Rab5. (**A-F**) Confocal images of NMJ synapses at muscle 4 at A2 hemisegment in *D42-Gal4*/*UAS-Tkv-EGFP* (A-C) and *D42-Gal4*, *AP2σ*^*KG02457*^/*AP2σ*^*ang7*^, *UAS-Tkv-EGFP* (D-F) labelled for Tkv (green) and early endosomal marker, Rab5 (magenta). Arrows in panel C show colocalization between Tkv-EGFP and Rab5 in control NMJ synapses, whereas the two signals are distinct at the mutant synapses (panel F). Scale bar in F represents 5 μm. (**G**) Histogram showing the percentage of Rab5 punctae positive for Tkv-EGFP in *D42-Gal4*/*UAS-Tkv-EGFP* (65.74 ± 3.75) and *D42-Gal4*, *AP2σ*^*KG02457*^/*AP2σ*^*ang7*^, *UAS-Tkv-EGFP* (27.4 ± 3.82) synapses. Error bar represents standard error of the mean (SEM); the statistical analysis was done using Student’s *t*-test. \*\*\**p*<0.001. (**H-M**) Confocal images of boutons from muscle 4 at A2 hemisegment in control and *AP2σ*^*KG02457*^/*AP2σ*^*ang7*^, immunolabelled for Rab5 (magenta) and HRP (green). Scale bar in M represents 5 μm. (**N**) Histogram showing the Rab5 level in control (100.0 ± 3.0) and *AP2σ*^*KG02457*^/*AP2σ*^*ang7*^ (97.25 ± 4.81) synapse. Error bar represents standard error of the mean (SEM); the statistical analysis was done using Student’s *t*-test. ns, not significant.

The decreased colocalization of Tkv with Rab5 could also be due to a reduced level of Rab5 itself in *σ2-adaptin* mutants. To test this possibility, we quantified the levels of synaptic Rab5 at the mutant synapses. We found that the synaptic levels of the early endosomal marker, Rab5 (*w*^*1118*^: 100.0 ± 3.0 vs. *AP2σ*^*KG02457*^/ *AP2σ*^*ang7*^: 97.25 ± 4.81) or late endosomal marker, Rab7 (*w*^*1118*^: 100.0 ± 5.71 vs. *AP2σ*^*KG02457*^/ *AP2σ*^*ang7*^: 115.6 ± 8.17) were not altered in *σ2-adaptin* mutant (Figure 6H-N and Supplemental Figure S2). These results suggest that decreased colocalization between Tkv-EGFP and Rab5 is primarily due to compromised internalization of Tkv receptors from the presynaptic membrane.

Synaptic proteins such as Nwk have been shown to associate with Rab11 and regulates BMP receptor recycling. Moreover, mutants that affect the recycling of BMP receptors show elevated pMAD levels and an increased number of boutons [11, 28, 53]. This prompted us to assess any possible defects in the recycling endosomes in *σ2-adaptin* mutants. In order to test the defect in recycling endosome trafficking, we stained NMJs with recycling endosome marker Rab11. Interestingly, we observed a drastic reduction in synaptic Rab11 levels (*w*^*1118*^: 100.0 ± 5.73 vs. *AP2σ*^*KG02457*^/ *AP2σ*^*ang7*^: 61.81 ± 7.10, p≤0.001) in *σ2-adaptin* mutant. (Figure 7A-I). Synaptic Rab11 levels were restored to control levels upon neuronal expression of a *σ2-adaptin* transgene in the *σ2-adaptin* mutant (*D42-Gal4*, *AP2σ*^*KG02457*^/ *UAS-AP2σ*, *AP2σ*^*ang7*^:112.9 ± 8.29) (Figure 7E-F and 7I). Consistent with these results, we found that downregulation of other AP2 subunits i.e. α-adaptin (*D42-Gal4*> α-adaptin^RNAi^:65.4 ± 3.31, p≤0.001); β_2_-adaptin (*D42-Gal4*> β_2_-adaptin^RNAi^:75.63 ± 2.55, p≤0.001) and μ_2_-adaptin (*D42-Gal4*> μ_2_-adaptin^RNAi^:78.30 ± 4.5, p≤0.001) in the motor neurons results in a significant decrease in Rab11 levels when compared with controls (*w*^*1118*^:100 ± 3.15) (Supplemental Figure S3). Taken together, these data indicate that: a) AP2 complex regulates recycling endosomes, and b) compromised endocytosis and defective recycling of the endocytosed Tkv receptors results in accumulation of Tkv receptors at the presynaptic membrane.

**Figure 7.**
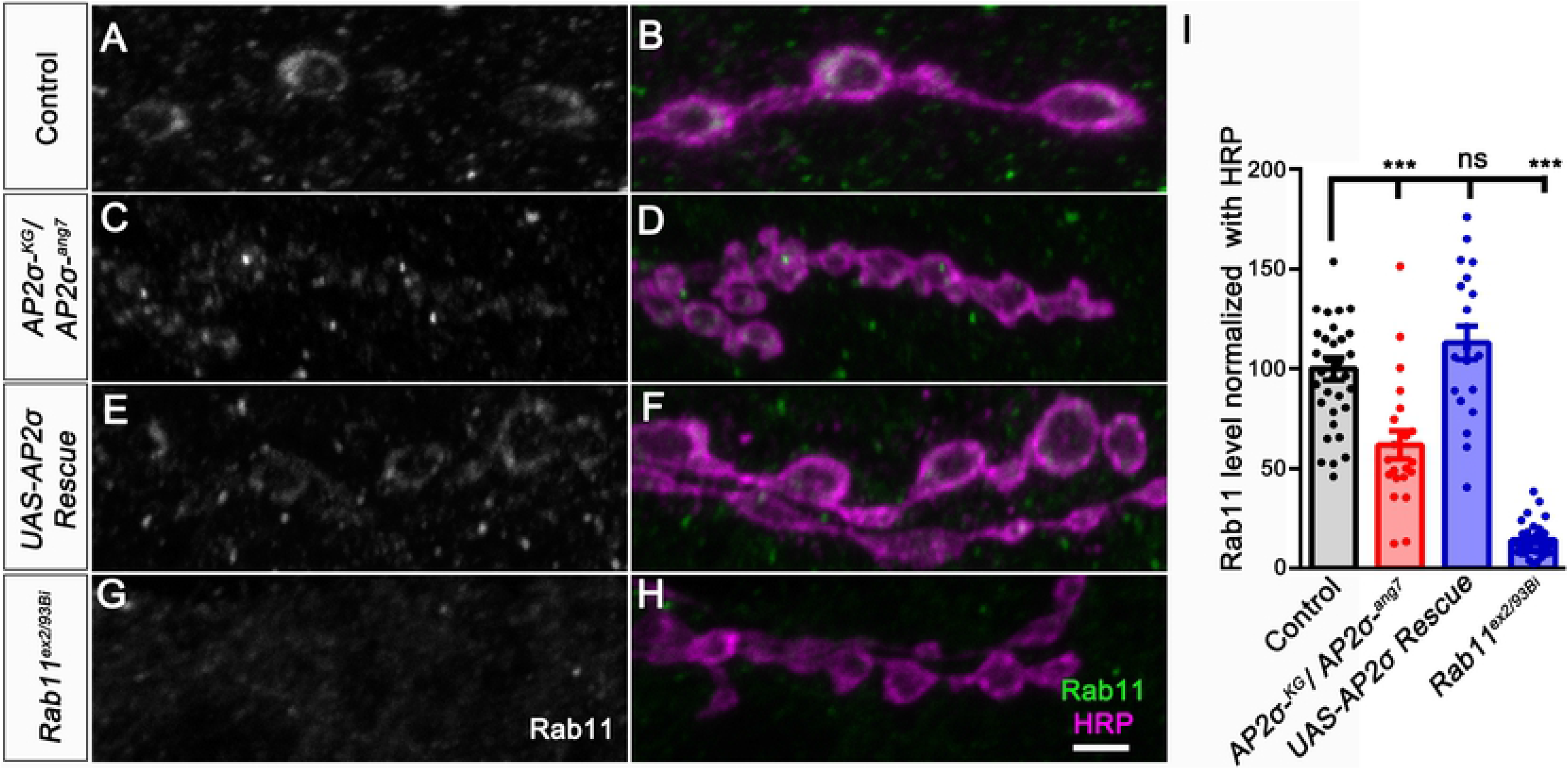
*σ2-adaptin* mutant synapses show a reduction in the recycling endosome marker, Rab11. (**A-H**) Confocal images of NMJ synapses at muscle 4 of A2 hemisegment in control (A-B), *AP2σ*^*KG02457*^*/AP2σ*^*ang7*^ (C-D), *D42-Gal4*, *AP2σ*^*KG02457*^/ *UAS-AP2σ*, *AP2σ*^*ang7*^ (E-F) and *Rab11*^*ex2/93Bi*^ (G-H) double immunolabeled with recycling endosomal marker, Rab11 (represented in grayscale/green) and neuronal membrane marker, HRP (magenta). Scale bar in (H) represents 3 μm. (**I**) Histogram showing the relative Tkv level normalized to HRP in control (100 ± 5.73), *AP2σ*^*KG02457*^/ *AP2σ*^*ang7*^ (61.81 ± 7.11); *D42-Gal4*, *AP2σ*^*KG02457*^/ *UAS-AP2σ*, *AP2σ*^*ang7*^ (112.9 ± 8.29) and *Rab11*^*EX2/93Bi*^ (14.21 ± 1.57) synapses. Error bar represents standard error of the mean (SEM); the statistical analysis was done using Student’s *t*-test. \*\*\**p*<0.001; ns, not significant.

### Rab11 mutants phenocopy the NMJ and BMP-signaling defects of *σ2-adaptin* mutants

Previous studies have shown that mutations in Rab11 results in NMJ morphological defects in *Drosophila* [28, 52]. Since *σ2-adaptin* functions in neurons to regulate BMP-signaling, we next asked if altering the levels of Rab11 specifically in neurons could phenocopy *σ2-adaptin* mutants and alter BMP signaling. Interestingly, we found that neuronal expression of a dominant-negative form of Rab11 (*Rab11*^*S25N*^) phenocopies the NMJ morphological defects of *σ2-adaptin* mutations and shows NMJ overgrowth (*w*^*1118*^: 1.56 ± 0.06), (*Rab11*^*ex2/93Bi*^:2.77 ± 0.11, p≤0.001), (*UAS-YFP-Rab11*^*S25N*^ /+; *D42-Gal4*/+: 2.26 ± 0.08, p≤0.001) and (*AP2σ*^*KG02457*^/ *AP2σ*^*ang7*^: 2.83 ± 0.12, p≤0.001) (Figure 8A-D and 8Q). Expressing wild type or constitutively active form of Rab11 does not alter the synaptic morphology (Supplemental Figure S4). Consistent with its role in BMP signaling and NMJ growth, *Rab11* mutants as well as animals expressing a dominant-negative form of Rab11 in motor neurons resulted in the accumulation of pMAD at the NMJ synapses (*w*^*1118*^: 100 ± 5.87), (*Rab11*^*ex2/93Bi*^: 147.9 ± 8.58, p≤0.01), (*UAS-YFP-Rab11*^*S25N*^ /+; *D42-Gal4/+:* 134.2 ± 4.36, p≤0.05) and (*AP2σ*^*KG02457*^/ *AP2σ*^*ang7*^: 218.8 ± 7.5, p≤0.001) (Figure 8E-L and 8R). We further assessed Tkv-EGFP levels at the Rab11 mutant synapses and found that *Rab11* mutants show increased Tkv levels at the synapse (*D42-Gal4, Rab11^ex2^/ Tkv-EGFP, Rab11^93Bi^*:189.5 ± 10.57) compared to control (*D42-Gal4/Tkv-EGFP*: 100 ± 8.71) (Figure 8M-P and 8S). This suggests that defective recycling of Tkv in *Rab11* mutant results in its accumulation at the synapse leading to increased BMP signaling and synaptic overgrowth. Interestingly, we also found that similar to *σ2-adaptin* mutants, downregulating neuronal levels of α-adaptin leads to reduced Rab11 and shows enrichment of pMAD (w^1118^:100 ± 6.66), (*D42-Gal4*> α-adaptin^RNAi^:278.7 ± 14.51, p≤0.001) and synaptic overgrowth (w^1118^:1.35 ± 0.064), (*D42-Gal4*> α-adaptin^RNAi^:2.59 ± 0.12, p≤0.001) at the *Drosophila* NMJ (Supplemental Figure S5). Taken together, these data suggest that Rab11-dependent trafficking of Tkv is deregulated in *σ2-adaptin* mutants leading to increased BMP signaling and synaptic overgrowth.

**Figure 8.**
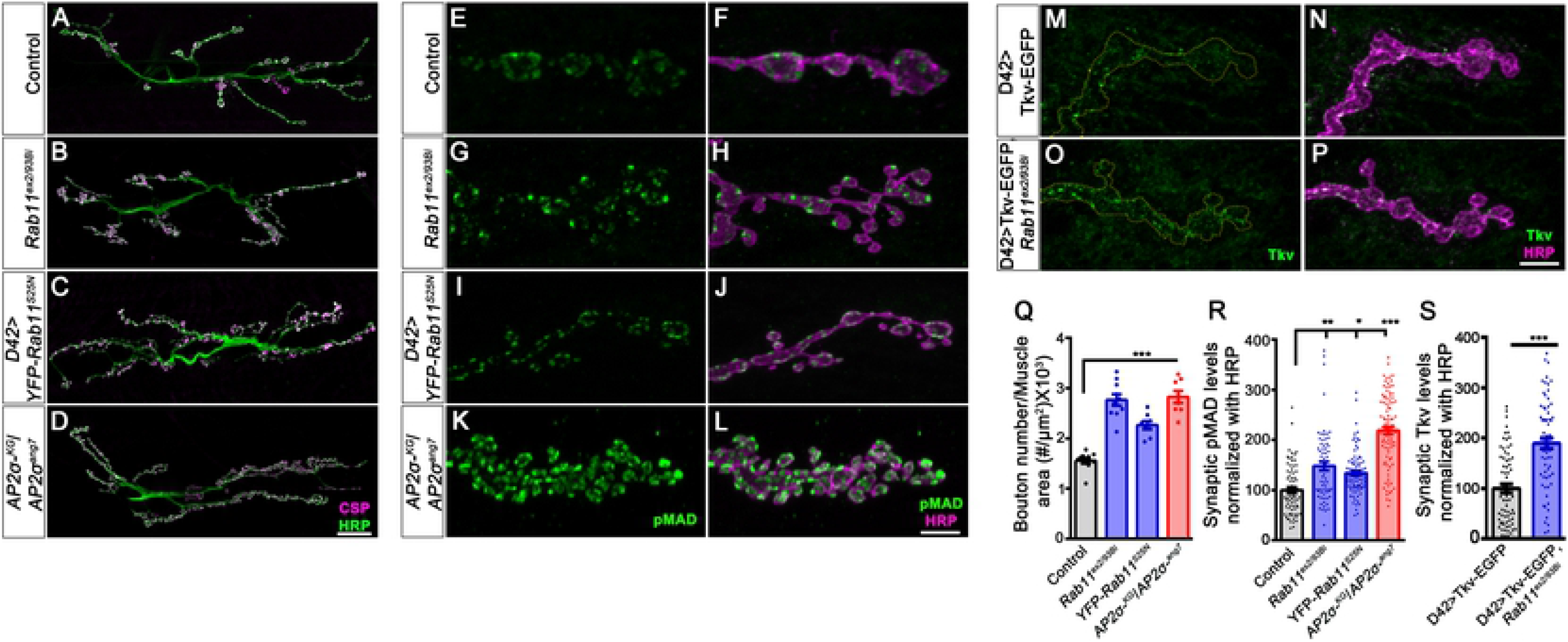
Rab11 mutants phenocopy *σ2-adaptin* mutations and show elevated levels of synaptic pMAD and Tkv receptors. (**A-D**) Confocal images of NMJ synapses at muscle 6/7 NMJ at A2 hemisegment in control (A), *Rab11*^*ex2/93Bi*^ (B), *D42-Gal4* driven dominant negative *YFP-Rab11*^*S25N*^ (C) and *AP2σ*^*KG02457*^*/AP2σ*^*ang7*^ (D) double immunolabeled with a pre-synaptic synaptic vesicle marker, CSP (magenta) and a neuronal membrane marker, HRP (green) to mark the bouton outline. Scale bar in D represents 20 μm. (**E-L**) Confocal images of NMJ from muscle 4 at A2 hemisegment in control (E-F), *Rab11*^*ex2/93Bi*^ (G-H), *D42-Gal4* driven dominant-negative *YFP-Rab11*^*S25N*^ (I-J) and *AP2σ*^*KG02457*^*/AP2σ*^*ang7*^ (K-L) double immunolabeled with pMAD (green) and a neuronal membrane marker, HRP (magenta) to mark the bouton outline. Scale bar in L represents 5 μm. (**M-P**) Confocal images of NMJ from muscle 4 at A2 hemisegment in control *D42-Gal4* driven Tkv-EGFP (M-N) and *D42-Gal4* driven Tkv-EGFP, *Rab11*^*ex2/93Bi*^ (O-P). Scale bar in P represents 5 μm. (**Q**) Histogram showing the average bouton number normalized to the muscle area from muscle 6/7 NMJ at A2 hemisegment in control *w*^*1118*^ (1.56 ± 0.06), *Rab11*^*ex2/93Bi*^ (2.77 ± 0.11), *D42-Gal4* driven dominant-negative *YFP-Rab11*^*S25N*^ (2.26 ± 0.08) and AP*2σ*^*KG02457*^/ AP2σ^*ang7*^ (2.83 ± 0.12). Error bar represents standard error of the mean (SEM); the statistical analysis was done using one-way ANOVA followed by post-hoc Tukey’s test. \*\*\**p*<0.001. (**R**) Histogram showing the levels of pMAD normalized with HRP from muscle 4 at A2 hemisegment in control (100 ± 5.87), *Rab11*^*ex2/93Bi*^ (147.9 ± 8.58), *D42* driven YFP-*Rab11*^*S25N*^ (134.2 ± 4.36), and *AP2σ*^*KG02457*^/ *AP2σ*^*ang7*^ (218.8 ± 7.5). Error bar represents standard error of the mean (SEM); the statistical analysis was done using one-way ANOVA followed by post-hoc Tukey’s test. \*\*\**p*<0.001, **p<0.01, *p<0.05. (**S**) Histogram showing the relative Tkv level normalized to HRP in *D42-Gal4* driven *Tkv-EGFP* (100 ± 8.71), and *D42-Gal4*, *Rab11*^*ex2*^/ Tkv-EGFP*, Rab11*^*93Bi*^ (189.5 ± 10.57). Error bar represents standard error of the mean (SEM); the statistical analysis was done using Student’s *t*-test. \*\*\**p*<0.001.

**Figure 9.**
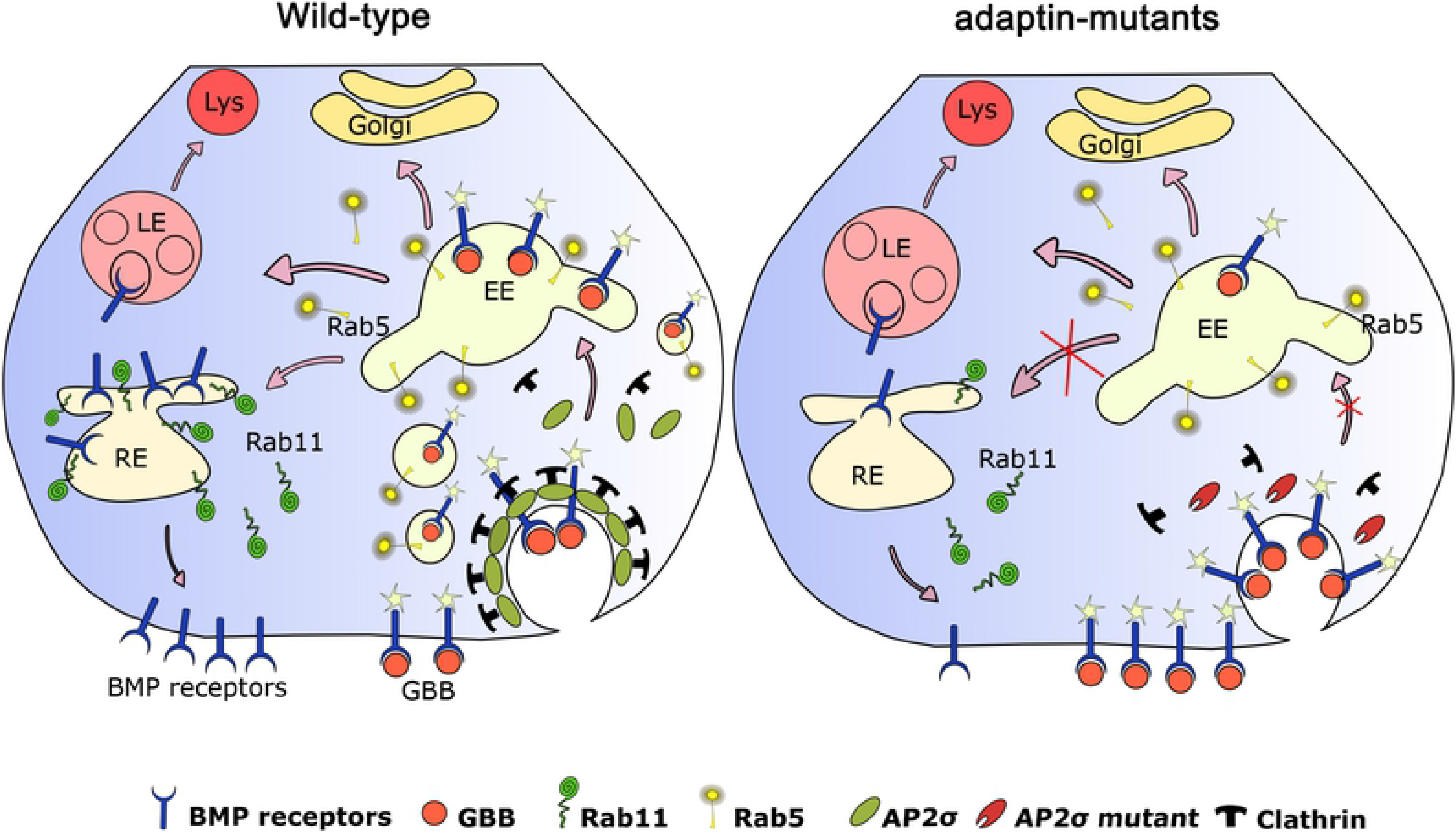
Model depicting the role of σ2-adaptin in BMP receptor trafficking at the NMJ. The model depicts a novel function of σ2-adaptin in BMP receptor trafficking at the *Drosophila* NMJ. Retrograde NMJ growth signaling in *Drosophila* involves Gbb ligand that is secreted from postsynaptic muscles and binds to BMP receptors on the presynaptic membrane to activate them [7, 12]. Activated receptors are then internalized through CME and fuse with early endosomes to trigger the downstream signaling cascade [71]. From early endosomes, receptors either get sorted to the Rab11-positive recycling endosomes that recycle them back to the presynaptic membrane or are sorted for lysosomal degradation [9, 28]. Depleting σ2-adaptin/AP2-complex perturbs endocytosis and Rab11-mediated recycling of type-I BMP receptor, Thickveins (Tkv) leading to its accumulation at the presynaptic membrane and early endosomes. Enrichment of Tkv receptors at the presynaptic membrane in the early endosome leads to elevated BMP signaling resulting in synaptic overgrowth.

## DISCUSSION

High fidelity neurotransmission depends on the proper endocytosis of synaptic vesicles after their fusion with the pre-synaptic membrane. Defects in endocytosis not only perturb synaptic transmission but also have been implicated in deregulated synaptic growth [3, 16, 54, 55]. The striking growth defects at the NMJ in *endo*, *synj* and *σ2-adaptin* mutants [3, 54–56] are a consequence of perturbing the endocytic machinery. However, our study on *σ2-adaptin* showed no change in levels of endocytic proteins like Endo, Synj, and Dynamin [8]. Therefore, we sought to address the role of CME in synaptic growth signaling. Defects in endocytosis, in general, have been linked to the BMP signaling cascade that drives the expression of growth-promoting genes through the activity of pMad [1, 8, 11, 15, 16]. The signaling activity, many a time, is dependent on intracellular traffic that is in part dependent on endocytosis of activated receptors, ultimately impinging on signaling pathways, particularly the BMP, JNK, and Wingless pathways [1, 4, 7, 8, 15, 57]. Here, we show for the first time a genetic interaction between σ2-adaptin and BMP pathway components. We further provide compelling evidence that the synaptic overgrowth phenotype in *σ2-adaptin* mutant is due to defective Rab11-mediated trafficking of type I BMP receptor, Thickveins.

### σ2-adaptin interacts with BMP pathway components to regulate neuronal BMP signaling pathway

Increasing evidence suggest the role of endocytosis in the regulation of synaptic signaling and NMJ growth. Endosomal trafficking of BMP receptors is a crucial regulatory feature that governs synaptic growth. Various proteins are known to interact with BMP receptors and either facilitate or impede the signaling cascade. A recent study in cultured mammalian cells reports one such protein, Angiomotin130 (AMOT130), with a coiled-coil motif and a C-terminal PDZ motif that interacts with the BMP receptor BMPR2 and facilitates BMP-SMAD signaling. AMOT130 localizes to endosomes and is thought to modulate GTPase signaling [58]. Endocytic proteins appear to be fascinating candidates as BMP receptor interactors.

Interestingly, *Drosophila* loss-of-function endocytic mutants correlate with elevated BMP signaling and neuronal overgrowth phenotype [9, 17, 20]. Consistent with this, we showed increased Tkv levels at the NMJ results in an elevated level of BMP pathway in *σ2-adaptin* mutants. If the BMP pathway is responsible for the synaptic overgrowth in *σ2-adaptin* mutants, then manipulating the levels of BMP signaling components should rescue the NMJ phenotype. In agreement, we show that by partially reducing BMP receptors Tkv, Wit, and cytosolic co-Smad molecule, Medea significantly rescues the phenotype. We further confirmed the interaction between σ2-adaptin and BMP signaling by showing that σ2-adaptin genetically interacts with the negative regulator of BMP signaling, the inhibitory Smad, Dad. Transheterozygotes of *dad* and *σ2-adaptin* mutants have an increased number of boutons compared to heterozygotes of either mutant alone.

### σ2-adaptin regulates trafficking of type I BMP receptor, Thickveins from the plasma membrane to early endosomes

BMP signaling has been extensively studied in the context of neuronal growth in which activated type I receptor, Tkv is endocytosed in the form of vesicles and fuse with early endosomes to activate downstream signaling molecules. The signaling is attenuated when these activated receptor-containing vesicles recycle back to the plasma membrane or fuse with lysosomes to be degraded [17, 59]. Trafficking of these receptors into and out of such endosomes provides an additional tier for spatial and temporal modulation of signal transduction. The members of the Rab family of small GTPases regulate various stages of endocytosis [24]. Our immunocytochemistry data show elevated Tkv receptor levels at the synapses and motor neuron soma (data not shown) of *σ2-adaptin* mutants. Besides, levels of Rab11 (known for its role in the recycling of Tkv receptor) is reduced by half in *σ2-adaptin* mutant synapses. Interestingly, levels of early and late endosomes marked with Rab5 and Rab7, respectively, remain unaffected at the *σ2-adaptin* mutant synapses with a much-reduced colocalization between Rab5 and Tkv receptor punctae compared to controls. Besides, the intensity profile of Tkv and HRP across the bouton shows that *σ2-adaptin* mutant has a higher intensity of Tkv at the membrane compared to control, indicating that a significant proportion of the Tkv receptors are accumulated at the presynaptic membrane

Tkv receptors could be accumulated either at the plasma membrane due to inefficient CME or at the early endosomes caused by inefficient recycling. If Tkv were accumulated at the early endosomes, we would expect greater colocalization with Rab5, which is not the case. However, based on reduced Rab11 staining at the mutant synapses, we conclude that the portion of the receptors that remain in Rab5 positive early endosomes fail to recycle back to the plasma membrane. This conclusion also fits with previous observations that defective CME results in the accumulation of endosome-like structures in cultured hippocampal neurons [30] and is substantiated by our electron microscopy data. The link between clathrin-mediated dynamin-dependent endocytosis and BMP signaling is still a contentious topic. A recent study using human umbilical vein endothelial cells (HUVECs) has shown that treating these cells with BMP-9 triggered caveolin-1 and dynamin-2 mediated endocytosis of its receptor, activin-like kinase 1 (ALK-1). Surprisingly, this ALK-1 endocytosis was not mediated by Clathrin heavy chain [60]. At the *Drosophila* NMJ, perturbing endocytosis results in upregulated BMP signaling [11], whereas in *Drosophila* wing discs and intestinal stem cells, endocytosis facilitates the signaling cascade by internalizing Tkv [61, 62], pointing towards a tissue-specific mechanism. Our results suggest a model where bulk membrane endocytosis is insufficient in removing Tkv from the plasma membrane; besides, the synapses in *σ2-adaptin* mutants fail to recycle remaining receptors from early endosomes leading to enhanced signaling and drastic NMJ growth defects.

### Functional and morphological aspects of σ2-adaptin-mediated BMP signaling can be delineated

Morphological features of synapses often dictate functional outcomes, and physiological analyses of BMP signaling mutants reveal the same. In BMP type II receptor mutant, *wit* larvae, the size of the NMJ is greatly reduced with concomitant reduced evoked excitatory potentials [10, 12]. Analyses of BMP type I receptor mutants, *tkv* and *sax*, co-Smad *medea,* and transcription factor *mad*, all have smaller synapses with severe functional deficits [6]. The same is true for the muscle-secreted BMP ligand, Gbb. *gbb* mutant larvae also exhibit shorter NMJs with severely reduced evoked potentials [7]. *σ2-adaptin* mutant synapses, however, show a modest reduction in evoked potentials, and the protein is dispensable for maintaining basal synaptic transmission [3]. Rundown of EJP amplitudes during high-frequency stimulation is used to measure endocytic defects. Synaptic mutants implicated in CME, such as *endophilin*, *synj* and *dap160*, show a rapid stimulus-dependent decline in EJP amplitude that recovers following a period of rest after the high-frequency stimulation paradigm [19, 55]. In our previous study, we had reported that *σ2-adaptin* mutants do not recover from synaptic depression even after a period of rest following cessation of high-frequency stimulation [3]. This observation suggested that in addition to its requirement in synaptic membrane retrieval, the σ2-adaptin function is also required during the much slower process of SV trafficking, possibly at one of the rate-limiting steps in SV regeneration. This result is now supported by our EM data that shows an accumulation of endosome-like structures at the mutant synapses. EJP and high-frequency recordings from *σ2-adaptin* mutant synapses with one copy of *tkv*^*7*^ did not show any rescue in synaptic function. These data drive the conclusion that partial reduction of BMP pathway components can only rescue morphological defects in *σ2-adaptin* mutants but not functional aspects and that morphological and functional deficits can be delineated in these mutants. Besides, a partial rescue of bouton size and bouton clustering at the NMJ argues for possible deregulation of multiple signaling pathways in σ2-adaptin mutants that remains to be explored.

Our study uncovers and extends the existing knowledge of synaptic growth signaling and endocytosis. We provide four lines of evidence on the critical role of σ2-adaptin in modulating BMP-dependent synaptic growth signaling at the *Drosophila* NMJ. First, we show using genetics that the morphological defects in *σ2-adaptin* mutant synapses can be partially rescued by introducing a mutant copy of the BMP receptors, *tkv,* and *wit*. We also show a direct epistatic interaction between σ2-adaptin and the inhibitory Smad, Dad. Second, using immunohistochemistry, we show that *σ2-adaptin* mutant synapses accumulate Tkv at the plasma membrane and some of these receptors that are endocytosed and make it to the early endosomes fail to recycle back to the plasma membrane due to decreased Rab11 GTPase. Third, our electrophysiology data establish that morphological and functional defects can be delineated in *σ2-adaptin* mutants. Finally, our electron micrographs provide conclusive evidence showing the presence of large endosomes that match with our speculation that σ2-adaptin is critically required at a later step of vesicle regeneration following endocytosis from the plasma membrane. Partial rescue of bouton size and bouton clustering argues for possible deregulation of multiple signaling pathways in σ2-adaptin mutants.

This study thus opens new avenues where the role of other CME components and their interaction with various growth signaling pathways can be studied. Since receptor localization and regulation appears to be the central theme in modulating BMP signaling and synapse growth, it will be interesting to perform structure-function analysis of BMP receptors and identify key residues/motifs that interact with AP2 and facilitate its endocytosis. Mutating tyrosine-based signal (YXXϕ) and dileucine-based signal ([DE]XXXL[LI]) motifs in Tkv and Wit could lead to further understanding of these intricate interactions.

## MATERIALS AND METHODS

### Fly stock

Flies were grown and maintained at 25°C temperature in a standard cornmeal medium as described in [3]. The wild type *w*^*1118*^ was used as control unless otherwise stated. Genetic combinations and recombinations were made using standard genetic crosses. All the mutants, controls, and rescued larvae were grown under the non-crowded condition on apple agar plates with yeast paste dollop. The following stocks were obtained from Bloomington *Drosophila* Stock Center (BDSC)-*tkv*^*7*^ (BL3242), *medea* (BL7340), *wit*^*A12*^(BL5173), *dad*^*j1E4*^ (BL10305), *AP2σ*^*KG02457*^ (BL13478), UAS-Rab11-GFP (BL8506), α-adaptin RNAi (BL32866), β2-adaptin RNAi (BL28328), μ2-adaptin RNAi (BL28040). Other lines used in this study are: *D42-Gal4* [63, 64], UAS-Tkv-EGFP (BL51653) [65], UAS-YFP-Rab11^S25N^ (BL9792) [66], UAS-Rab11^Q70L^-GFP (BL23260) [67], Rab11^EX2^ and Rab11^93Bi^ [52].

### Antibodies and Immunocytochemistry

Wandering third instar larvae were dissected in cold calcium-free HL3 saline (70 mM NaCl, 5 mM KCl, 20 mM MgCl2, 10 mM NaHCO3, 5 mM Trehalose, 115 mM sucrose, and 5 mM HEPES, pH 7.2) to expose the NMJs and fixed in 4% paraformaldehyde in phosphate-buffered saline (PBS, pH 7.2) for 30 min at room temperature. Fillets were then washed in PBS containing 0.15% Triton X-100, blocked for 1 hour with 5% bovine serum albumin (BSA) followed by overnight incubation with primary antibody at 4°C. The monoclonal antibody anti-CSP (1:100) was obtained from the Developmental Studies Hybridoma Bank (DSHB). The polyclonal antibody against Rab5 [68] was a gift from Marino Zerial, Max Planck Institute, Germany. The polyclonal Rab7 and Rab11 [44, 69] antibodies were a gift from Tsubaka Tanaka, RIKEN Center for Developmental Biology, Japan. The secondary antibodies conjugated to Alexa Fluor 488 and Alexa Fluor 568 (Molecular Probes, Thermo Fisher Scientific) were used at 1:800 dilution. The Alexa Fluor 488 or Rhodamine conjugated anti-HRP (Jackson Immunoresearch, USA) were used at 1:800 dilution. Stained larval fillets were mounted in VECTASHIELD (Vector Laboratories, Burlingame, CA). All the images were captured with a laser scanning confocal microscope (LSM780, Carl Zeiss, Jena Germany or FV3000, Olympus Corporation, Japan).

### Electrophysiology

All the intracellular recordings were performed on wandering third instar larvae as described previously [3]. Briefly, HL3 buffer containing 1.5 mM Ca^2+^ was used for the larval dissection. Recordings from muscle 6 of A2 hemisegment were performed using sharp glass electrodes having a resistance of 20-25 MΩ resistance. Miniature EJPs (mEJPs) were recorded for 60 seconds, followed by recordings of EJPs at 1 Hz stimulation. For High-frequency recording, nerves were stimulated at 10 Hz, and EJPs were recorded for 5 minutes. For recording EJPs, stimulation pulse was delivered using Grass S88 stimulator (Grass Instruments, Astro-Med, Inc). The signals were amplified using Axoclamp 900A, digitized using Digidata 1440A, and acquired using pClamp10 software (Axon Instruments, Molecular device, USA). Muscles with resting membrane potential between −60 mV and −75 mV were used for analysis. The data were analyzed using the Mini Analysis program (Synaptosoft, Decatur, USA).

### Colocalization and Intensity profile

Confocal images of muscle 4 NMJ at A2 hemisegment were used to quantify the colocalization percentage between Rab5 and Tkv. From each NMJ, around ten random puncta were chosen and analyzed manually. To analyze the colocalization using motion tracker software, ROI was created around the NMJ. The intensity and size threshold were optimized for each NMJ so that the software randomly selects at least 10-15 puncta in each NMJ for the colocalization analysis. To plot the intensity profile, a single bouton section was used, and the intensity of Tkv and HRP was analyzed throughout a line using Fiji/ ImageJ software. The graph was plotted in the excel file using the intensity values obtained from Fiji/ImageJ software. As the intensity of Tkv in control was too less to plot the graph, all the intensity values were multiplied by two.

### Electron Microscopy

TEM was performed as described previously [70]. Third instar larvae were dissected in cold PBS. The larval fillets were then fixed in 0.12M cacodylate buffer containing 2% glutaraldehyde for 10 minutes at room temperature, transferred to a fresh fixative, and kept overnight at 4°C. The fillets were postfixed for 1 hour with 2% osmium tetroxide (OsO4) solution prepared in 0.12M cacodylate buffer. The samples were rinsed with 0.12M cacodylate buffer followed by washes with distilled water to avoid precipitation of cacodylate with Uranyl acetate. Subsequently, the samples were subjected to en bloc staining with 2% uranyl acetate. The stained fillets were again washed with distilled water and dehydrated using graded solutions of ethanol before final infiltration of the samples through propylene oxide for 30 minutes. Stained and dehydrated fillets were embedded in epoxy resin and hardened overnight at 60°C. Muscles embedded in epoxy resin were sectioned at 60 nm. Ultrathin sections of the muscles stained with 2% uranyl acetate (in 70% ethanol) and 1% aqueous lead citrate were examined at 120 KV on a Tecnai G2 Spirit BioTWIN (FEI, USA) electron microscope. The number of synaptic vesicles per bouton were counted manually using the Multi-point tool in ImageJ/Fiji software and then divided by their respective bouton areas to obtain the vesicle density /μm^2^ area of a bouton. For vesicle size, diameters of at least 100 vesicles from 10 bouton sections of each genotype were used for quantification.

### Quantification and statistical analysis

For fluorescence quantification, images were captured using a laser scanning confocal microscope (LSM780; Carl Zeiss or FV3000, Olympus). All the control and experimental fillets were processed in the same way, and the fluorescence images were captured under the same settings for every experimental set. For bouton quantification, CSP labeled structures were counted at muscle 6/7 of A2 hemi-segment. The number of boutons from each NMJ was normalized to the respective muscle area. To calculate the bouton number, NMJs from A2 hemisegment were captured using a 40x objective, and all the CSP positive boutons were counted manually in ImageJ/Fiji software. For muscle area quantification, images from A2 hemisegment were captured using 20x objective, and the area was quantified using ZEN2 software (Carl Zeiss, Germany). For bouton number quantification, the total number of boutons per NMJ were divided by their respective muscle area. For fluorescence intensity quantification, NMJs from muscle 4 were captured using a 60X objective. For each NMJ, the fluorescence intensity from each bouton was subtracted from the background intensity, and the average intensity was normalized to the control. The fluorescence intensity was calculated using ImageJ/Fiji software. For bouton area quantification, NMJs from muscle 6/7 at A2 hemisegment were captured, and area was calculated by drawing a free-hand sketch around CSP positive bouton using ImageJ/Fiji software. The number of samples used for analysis is shown in the respective figure’s histogram or mentioned in the legend. For multiple comparisons, one-way ANOVA followed by Post-hoc Tukey’s test, and Student’s t-test was used. GraphPad Prism 8 was used to plot the graph. Error bars in all the histograms represent the standard error of the mean (SEM). \**P*<0.5, \*\**P*<0.01, \*\*\**P*<0.001.

## Acknowledgements

We thank Dr. Tsubasa Tanaka for sharing Rab5, Rab7, and Rab11 antibodies and Dr. Marino Zerial for Rab5 antibodies. We thank Dr. Avital Rodal for sharing Rab11 mutants. We thank the Bloomington Drosophila Stock Center (BDSC), Vienna Drosophila RNAi Centre (VDRC), and Drosophila Genomics Resource Centre (DGRC) for fly stocks and Developmental Studies Hybridoma Bank (DSHB), the University of Iowa for monoclonal antibodies. We thank Manish Jaiswal and Jeet Kalia for helpful comments on the manuscript.

## Author Contribution

S.D.C., M.K.D., and V.K. conceived and designed the experiments. M.K.D., S.D.C., A.P., and S.M. performed the experiments. M.K.D., S.D.C., and V.K. analyzed and wrote the manuscript with inputs from other authors. R.P provided resources and guidance for the TEM experiments. The authors declare no competing or financial interests. All the authors have read and approved the final version of the manuscript.

